# Differential acquisition of cocaine and heroin self-administration in a rat model of internalizing versus externalizing temperament

**DOI:** 10.64898/2026.04.09.717453

**Authors:** M.A. Emery, A. Parsegian, S. Koonse, E.K. Hebda-Bauer, K. Lee, B.D. Luma, S.E. Chang, J. Becker, S.B. Flagel, S.J. Watson, H. Akil

**Affiliations:** Michigan Neuroscience Institute, University of Michigan.; Department of Psychology, University of Michigan.; Department of Psychiatry, Michigan Medicine, University of Michigan

## Abstract

Substance Use Disorders (SUDs) constitute a major and rising public health concern. In addition, there is a growing appreciation that different classes of addictive substances are likely to lead to qualitatively different types of SUDs requiring differing treatment and relapse prevention strategies to be most effectively managed. Biological temperament, particularly on the internalizing – externalizing axis, is well established to influence addiction susceptibility. Externalizing behavior has long been understood to predispose individuals to addiction through novelty-seeking, sensation-seeking and impulsivity, while internalizing behavior provides an alternate pathway into addiction via increased occurrence of comorbid disorders (anxiety, depression). Here, we utilize a selectively bred rat model of internalizing vs externalizing temperament (bred High Responders, representing genetically mediated externalizing behavior and bred Low Responders, representing internalizing behavior) to examine differences in the acquisition of self-administration of the prototypical psychostimulant cocaine and the prototypical opioid heroin (diacetylmorphine). We found that, as predicted, cocaine and heroin drove different patterns of acquisition in the two different bred lines of rats. Further, this was influenced by temperament in complex ways. Notably, in females the “telescoping effect” for opioid addiction-like behavior was primarily specific to externalizing temperament. These findings highlight the impact and interaction of many factors, including drug class, temperament, and sex, on the acquisition of drug-taking behavior. Additionally, these findings indicate that sex differences in addiction vulnerability may be influenced in part by biological temperament.

---

Opioid Use Disorder constitutes a significant public health challenge in the United States currently, with drug overdose deaths rising fivefold over the last two decades since 2001, and opioids constitute the lion’s share of those deaths (Spencer 2022). However, only a small proportion of those who use opioids, approximately a quarter to a third, will go on to develop problematic use that meets the criteria of a Substance Use Disorder (Anthony et al. 1994). For cocaine users, only 1 in 6 meet the criteria (Anthony et al. 1994). More recent research has found this risk may be considerably lower, due to under-reporting of recreational and occasional cocaine use, and nearer to 5-6% of all users (Wagner and Anthony 2002, O’Brien and Anthony 2005). Although many factors have been established to contribute to the risk of developing SUD, including genetics, socioeconomic status, home environment, stress, and others, there is currently no reliable way to predict who will go on to develop SUD following exposure to addictive substances.

Multiple animal models demonstrate that externalizing temperamental traits, such as novelty-seeking, sensation-seeking, and high impulsivity, represent significant baseline risk for developing pathological drug seeking and taking behavior. This reflects what is seen in human personality factors research, where multiple models identify substance use problems as a facet of externalizing temperament (Krueger 1999, Hopwood et al. 2008), and externalizing temperament and problematic substance use appear to partially share a genetic basis (Iacono et al. 2008).

One animal model developed to investigate this relationship is the High Responder/Low Responder to a Novel Environment (HR/LR) rat model, first described by Piazza et al. (1989), which demonstrated that a high degree of locomotor response to a novel environment (HR) was reliably predictive of increased levels of psychostimulant self-administration behavior (Piazza et al. 1989, Piazza et al. 1990, Piazza et al. 1991) and locomotor response and sensitization (Hooks et al. 1991) compared to animals exhibiting a low degree of locomotor response to a novel environment (LR); a phenomenon that has been extended beyond psychostimulants to several other drugs of abuse, including nicotine (Suto et al. 2001), opioids (Ambrosio et al. 1995) and alcohol (Nadal et al. 2002). Later work demonstrated that the HR phenotype was also predictive of externalizing behavior styles across a range of behaviors, while LRs demonstrate internalizing behavior styles (Dellu et al. 1996, Kabbaj et al. 2000, Mällo et al. 2007, Jama et al. 2008). Our (Akil) lab has used a selective breeding strategy to develop rat lines selectively bred for high response to novelty (bHR) or low response to novelty (bLR), to investigate the heritable nature of this trait and its genetic underpinnings more systematically (Stead et al. 2006, Birt et al. 2021). The bHR/bLR lines have reconfirmed and expanded upon these findings, underscoring the broader ‘externalizing vs internalizing’ nature of the model. Indeed, Piazza’s HR/LR model, as a model of drug addiction behavior and genetics, is now understood to be predicated on the relationship between externalizing behavior and basal propensity to take drugs (both observed to greater degree in HRs) (Flagel et al. 2010a).

However, the LR phenotype (and internalizing temperament more generally) should not be considered to conversely represent a lower propensity or ‘resilient’ phenotype regarding substance use and addiction. It has been observed in humans that internalizing behavior represents not a resiliency to drug taking but rather an alternate pathway to drug abuse (Hussong et al. 2011), with individuals above average on either externalizing or internalizing tendencies having greater risk than median responders to abuse a wide range of substances including alcohol, opiates, and stimulants (Le Bon et al. 2004). Nearly a quarter of those diagnosed with a substance use disorder who are using illicit drugs (versus legal substances such as alcohol) have a comorbid diagnosis of anxiety, and 21% have comorbid depression (Conway et al. 2006). Nearly half of those abusing alcohol have either anxiety or depression (Alegría et al. 2010, Vasilieva et al. 2022), demonstrating the high degree of overlap between clinically significant internalizing behaviors and substance use. This seems to be reflected in the HR/LR model as well. Indeed, while both outbred LR and bLR rats demonstrate lower propensity to acquire psychostimulant self-administration at baseline (Piazza et al. 1989, Piazza et al. 1990, Piazza et al. 1991, Piazza et al. 2000, Davis et al. 2008, Cummings et al. 2011, Flagel et al. 2016), the experience of stress, particularly but not limited to social stress, is capable of shifting the LR phenotype into HR-like drug taking (Kabbaj et al. 2001) and behavioral sensitization (Aydin et al. 2021).

While the phenomenon of drug preference among addicts has long been noted anecdotally (Burroughs 1957), the concept has historically been under-explored in the scientific literature. There has been some indication in humans that drug preference is related to psychological traits that can be characterized on the externalizing/internalizing spectrum (Carrol and Zuckerman 1977, Flynn et al. 1995, Feldman et al. 2011), but the role of drug preference in drug choice is often overshadowed by drug availability. However, it is possible that the LR rat represents a genetic propensity not only for drug taking following stress, but a basal preference for a different drug experience. It was recently observed that outbred LR rats do not differ from outbred HRs in any examined facet of self-administration of the opioid remifentanil (Chang et al. 2022), despite strong differences in stimulant self-administration which is characteristic of the model (Piazza et al. 1989, Piazza et al. 1990, Piazza et al. 1991, Piazza et al. 2000). This may be indicative of differing drug preference between the lines, where internalizing LRs ‘dislike’ stimulants but readily take opiates, whereas sensation-seeking HRs have a higher basal preference for stimulants.

This reflects a relationship between personality and drug preference observed in humans but under-explored in the literature (Carrol and Zuckerman 1977). Despite multiple studies examining stimulant self-administration behavior, self-administration of opiates has not previously been examined in the selectively bred bHR/bLR model. Here, we examined the acquisition of heroin self-administration in male and female bHR and bLR rats, compared to the acquisition of cocaine self-administration, and found that the acquisition of heroin taking differs from the acquisition of cocaine taking in both lines.

## MATERIALS AND METHODS

### Animals

A total of 101 male and female Sprague–Dawley rats from our bred HR/LR colony (50 bHR and 51 bLR) from generations F70, 74, and 78 were used in this study. Rats were housed singly following surgery in standard controlled environmental conditions (22°C, 70% humidity, and 12-h light/dark cycle (males) or 14-h light-dark cycle (females), lights on at 05:00 (F) or 07:00 (M)) with food and water available *ad libitum*. All animal procedures were performed in accordance with the University of Michigan animal care committee’s regulations and followed the Guide for the Care and Use of Laboratory Animals: Eighth Edition (Council 2011).

### Surgical Procedures and Post-Operative Care

Rats were anesthetized using inhaled isoflurane (3-4% for induction, 2% ± 1% for maintenance of anesthesia). Indwelling jugular catheters were permanently implanted in the right jugular vein and threaded subcutaneously to a port emerging on the animal’s back at the midline approximately 1 cm caudal to the shoulder blades (Krueger et al. 2021). Postoperative pain was managed for a minimum of 48 hours and thereafter as necessary with NSAID drugs (flunixin meglumine, 5 mg/kg s.c.). Catheters were flushed upon implantation and then daily beginning 48 hours post-surgery with heparin (10 units) and gentamicin sulfate (0.1 mg) irrespective of body weight, throughout the duration of the experiment. Catheter patency was tested prior to beginning intravenous self-administration and weekly thereafter by administering 1 mg methohexital sodium (Par Pharmaceutical) in sterile saline via the catheter and observing for rapid onset of anesthesia. Any animal failing catheter patency at any point was removed from analysis from that point forward.

### Food Self-Administration Training

Beginning day 14 post-surgery, rats were trained to nose-poke for a 45-mg banana-flavored pellet (Bioserv; Product # F0059) using the same positioning of active and inactive noseports, and availability cues, as would later be utilized for drug self-administration. Rats could nose-poke on a fixed ratio (FR)-1 reinforcement schedule to receive up to 25 pellets per 1-hour session, one session per day for 5 days. Rats were not required to meet any specific criteria during this phase to advance to drug self-administration, but the acquisition of food self-administration was recorded for each rat.

### Drug Self-Administration

Following food self-administration, rats were allowed to self-administer cocaine hydrochloride (0.3 mg/kg/infusion) or heroin (diacetylmorphine hydrochloride, 0.02 mg/kg/infusion). Group sizes are summarized in Table 1. Start of a session was signaled by the illumination of the house light. Nosepokes in the active port (as trained during food SA) were reinforced by the extinguishing of the house-light, illumination of a cue light within the noseport, and the delivery of a 50 µL intravenous infusion delivered over ∼3 seconds. Following each infusion, a 20-second timeout period began, indicated by the continuing absence of the house-light and the extinguishing of the active noseport light, wherein additional nosepokes (at either port) were recorded but were without consequence. At the end of the 20s timeout period, the house-light would illuminate, signaling availability of drug, and the next nose-poke in the active port would be reinforced, beginning the timeout phase again. Rats were allowed to freely self-administer over a 2-hour session, capped at a maximum of 200 possible infusions to minimize overdose risk. Most rats never met this maximum for either drug. Drug self-administration proceeded M-F for 3 weeks, for a total of 15 sessions, with 48-h forced abstinence periods between D5-6 and D10-11. The general experimental timeline is presented in Figure 1.

**Figure 1.**
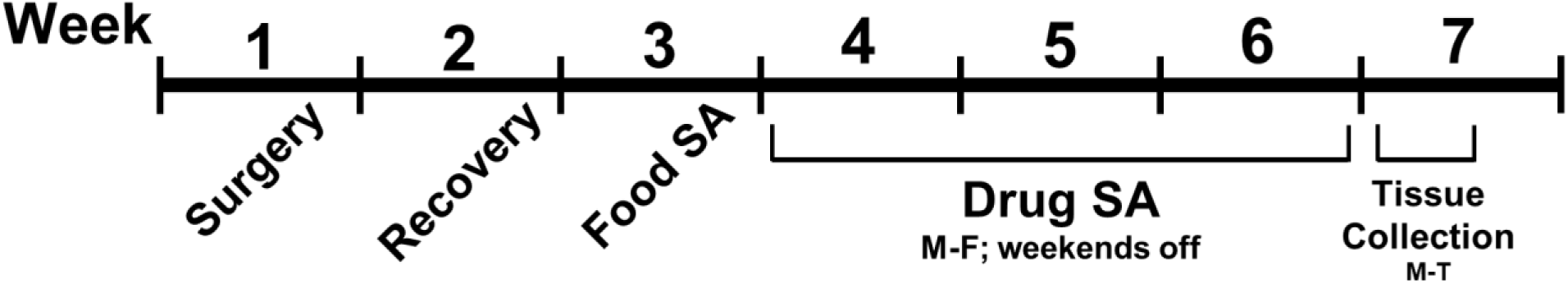
Schematic timeline of the experimental protocol. Rats underwent jugular catheterization surgery at approximately age P65-P75, 2 weeks following phenotype characterization by locomotor response in a novel environment. Following surgery, rats were given ∼1 week to recover, and then began food training for a week. Following food training, animals proceeded to drug self-administration for the next 3 weeks. Rats were selected to represent phenotypically ideal locomotor scores, and from as many different families as possible, to maximize genetic diversity. All rats were single-housed following surgery through the completion of the experiment.

**Table 1.**
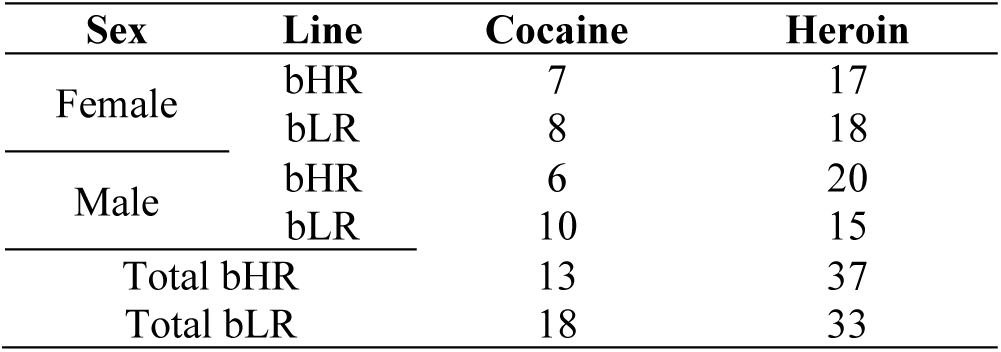
Summary of group sizes.

### Analysis

Mixed linear multilevel modeling was used to analyze the number of infusions taken, active nosepokes (an analog of drug seeking behavior, independent of reward availability), inactive nosepokes, and the average interinfusion interval (III), analyzing each drug independently (cocaine vs. heroin), with factors of line (bHR vs. bLR), sex (male vs. female) and time (average first 3 days vs. average last 3 days). In cases where sex was not determined to be a significant factor, as a main effect or in interaction with any other factor, it was dropped out as a variable (data for both sexes were pooled within line) and the model was re-run to increase statistical power. Post-hoc t-tests were run in cases where a significant interaction was found by the model, to determine the nature of the interaction. Analysis of coefficients of variation was performed using the Feltz-Miller asymptotic χ^2^ test (Feltz and Miller 1996).

## RESULTS

### Infusions

#### Rats modeling externalizing behavior self-administer more drug, regardless of drug class

For cocaine infusions taken, there was a significant main effect of line (*F*_1, 172_ = 40.94, *p* < 0.001), with bHRs taking significantly more infusions than bLRs (bHR mean ± SEM = 58.13 ± 4.15; bLR mean ± SEM = 23.47 ± 3.49, Figure 2a). Neither sex nor time were a significant factor, either as a main effect or in interaction with each other or line. See Table 1 for sample sizes.

**Figure 2.**
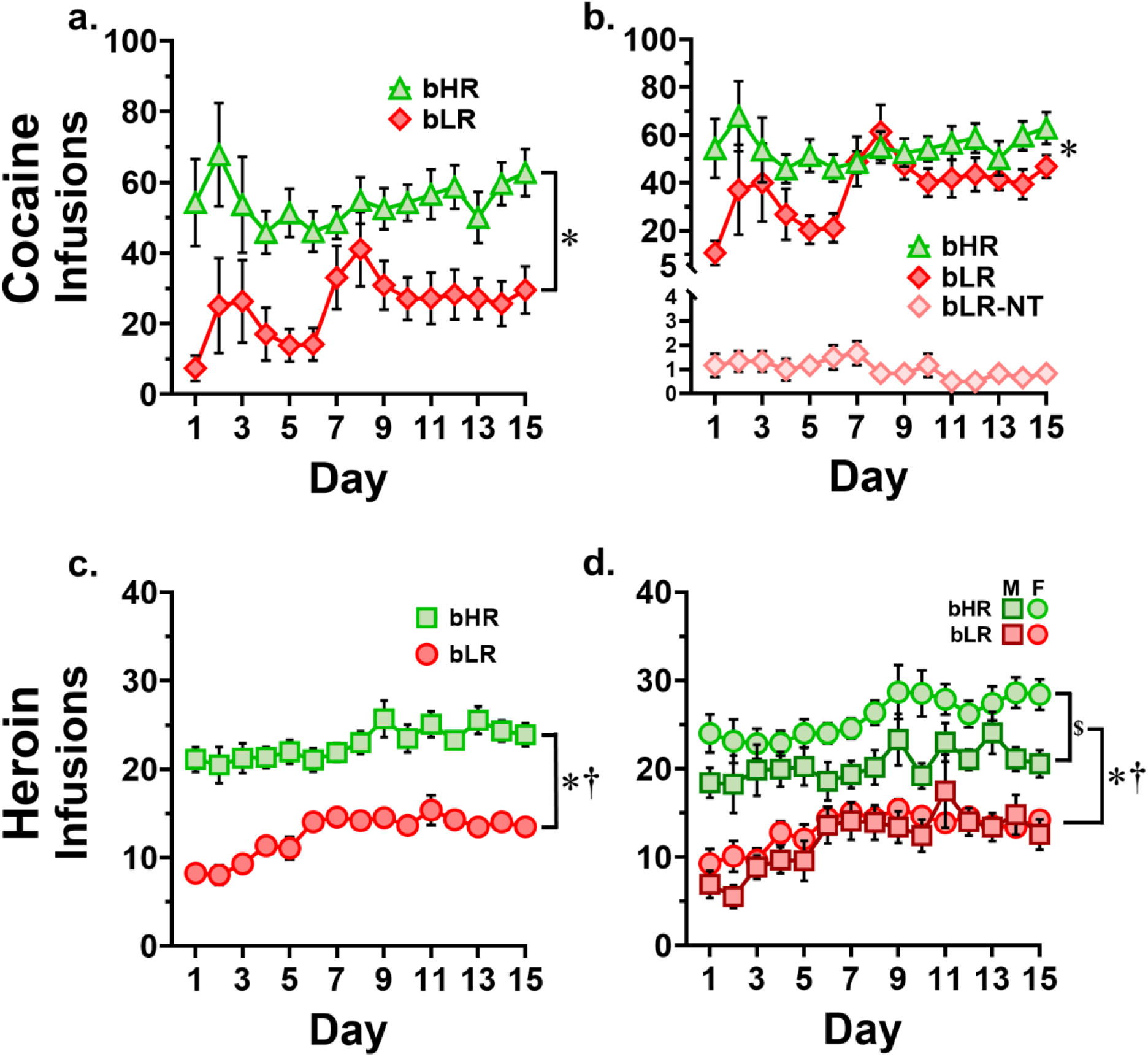
Drug infusions by day. **a.** Cocaine infusions, bHR vs. bLR. * main effect of line (*p* < 0.001). **b.** Cocaine, with bLR-non-takers (bLR-NT) displayed separately. * main effect of line (*p* < 0.001). **c.** Heroin infusions, bHR vs. bLR. * main effect of line (*p* < 0.001), † main effect of time (*p* < 0.001). **d.** Heroin, with sexes displayed separately. * main effect of line (*p* < 0.001), † main effect of time (*p* < 0.001), $ sex difference (*p* < 0.001). Note different y-axis scales.

For heroin infusions taken, there was a significant main effect of line (*F*_1, 402_ = 230.72, *p* < 0.001), with bHRs taking significantly more infusions than bLRs (bHR mean ± SEM = 22.76 ± 0.53; bLR mean ± SEM = 11.10 ± 0.55), as well as a significant main effect of time (*F*_1, 402_ = 32.69, *p* < 0.001), with both lines accelerating heroin taking levels over the course of the experiment (Figure 2c). See Table 1 for sample sizes.

#### Drug-specific differences in drug consumption patterns are phenotype-dependent

##### A strongly cocaine addiction-resistant subpopulation emerges only in the internalizing phenotype

Exploration of the data uncovered two distinct subpopulations among the bLR group in cocaine-taking behavior. By the second week of cocaine self-administration, two distinct subgroups have emerged. Most bLRs begin to accelerate cocaine taking, ultimately reaching consumption levels nearly that of bHRs (bLR infusions in the final three days, mean ± SEM = 42.4 ± 6.54; bHRs mean ± SEM = 57.89 ± 6.26). Despite cocaine taking levels approaching that of bHRs, a line difference still persisted between bHRs and the subgroup of bLRs who developed cocaine taking at the end of the experiment (average infusions across final 3 days, *t*_65_ = 3.24, *p* < .001). However, a smaller group of bLRs whom we have named “non-takers” remain at very low levels of cocaine taking (infusions, mean ± SEM = 1.03 ± 0.91) throughout the full 15 days of the experiment (Figure 2b).

##### Increased heroin taking in females emerges only in the externalizing phenotype

Exploration of the data also revealed a line-specific sex difference in self-administration behavior that is specific to heroin (Figure 2d). When sex was included as a variable for analysis of heroin infusions taken, there was a significant main effect of sex (*F*_1, 398_ = 20.1, *p* < 0.001), and a significant line by sex interaction (*F*_1, 398_ = 7.244, *p* = 0.007), in addition to the previously determined significant main effects of line and time. When given the opportunity to self-administer heroin, a line by sex interaction is driven by bHR females taking significantly more heroin than their male counterparts (average infusions across experiment, t_208_ = 4.155, *p* < 0.001); no sex difference exists in bLRs (t_182_ = 0.187, *p* = 0.852), indicating that the effect in bHRs is strong enough to drive the main effect. These data are summarized in Table 2.

**Table 2.** Summary of sex by line differences observed in self-administration behavior.

| Line | Sex | Cocaine |  | Heroin |  |
| --- | --- | --- | --- | --- | --- |
|  |  | Infusions | Active Nosepokes | Infusions | Active Nosepokes |
| bHR | Male | 55.78 (7.56) | 61.1 (7.0) | 20.39 (0.92) | 26.5 (1.1) |
|  | Female | 60.43 (3.82) | 75.2 (5.5) | 25.59 (0.81) | 36.6 (1.1) |
| bLR | Male | 19.57 (3.83) | 18.5 (3.8) | 10.36 (0.73) | 15.6 (1.1) |
|  | Female | 28.63 (6.28) | 23.2 (3.6) | 11.70 (0.55) | 17.8 (0.9) |

Data are presented as mean (SEM).

#### Drug-taking behavior in both lines is more variable for cocaine than for heroin

For both bHRs and bLRs, across the duration of the experiment, the degree of variation around the mean number of infusions, as determined by the coefficient of variation, is higher for cocaine than for heroin (bHRs *χ*^2^_1_ = 17.23, *p* < 0.001; bLRs *χ*^2^_1_ = 112.31, *p* < 0.001). Note that for cocaine infusions, this data was filtered to remove non-takers, as they represent an artificially high degree of variation from the mean score if left in. Interestingly, despite no significant sex difference in cocaine taking behavior, male bHRs show a higher degree of variation in cocaine taking than their female counterparts (*χ*^2^_1_ = 44.17, *p* < 0.001); this is not seen in bLRs (*χ*^2^_1_ = 0.95, *p* = 0.33). These data are summarized in Table 3.

**Table 3.**
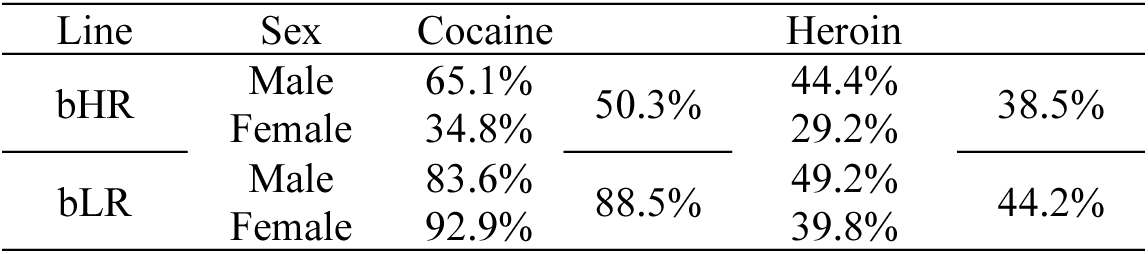
Summary of coefficients of variation of infusions.

| Line | Sex | Cocaine |  | Heroin |  |
| --- | --- | --- | --- | --- | --- |
| bHR | Male | 65.1% | 50.3% | 44.4% | 38.5% |
|  | Female | 34.8% |  | 29.2% |  |
| bLR | Male | 83.6% | 88.5% | 49.2% | 44.2% |
|  | Female | 92.9% |  | 39.8% |  |

### Active Nosepokes

Because active nosepokes occur in the absence of reinforcement and are theoretically limited only by the animal’s physical movement speed, it is possible for exceedingly high rates/levels of active nosepokes to be achieved which are more likely representative of a learned repetitive behavior or a drug-induced stereotypy than volitional action. For this reason, the Tukey Method was used to identify statistical outliers in active nosepoke data; those outliers were removed prior to analysis (Supplemental Figure 1). Although outliers were identified and removed individually by day, the identified outlying scores were associated with ‘repeat offender’ rats approximately half the time, further reinforcing the possibility of behavioral stereotypy. Data presented in Figure 3a includes all collected datapoints. Data presented in Figure 3b and analyzed here, is processed to remove outliers. Box-and-Whisker plots using the Tukey method and used to identify outliers for removal can be found in Supplemental Figure 1. For cocaine, there was a significant interaction of line by time (*F*_1, 118_ = 9.44, *p* = 0.003), and an interaction of line by sex (F = 4.56, *p* = 0.035) as well as main effects of both line (*F*_1, 118_ = 44.61, *p* < 0.001) and time (*F*_1, 118_ = 15.88, *p* < 0.001). Drug seeking behavior is higher in bHRs, but the main effect of time is driven entirely by the strength of the effect in bLRs, with bHRs not significantly increasing active nosepokes over time (average infusions, first 3 days vs last 3 days, *t*_66_ = 0.398, *p* = 0.692). In addition, the line difference is more pronounced in females than in males.

**Figure 3.**
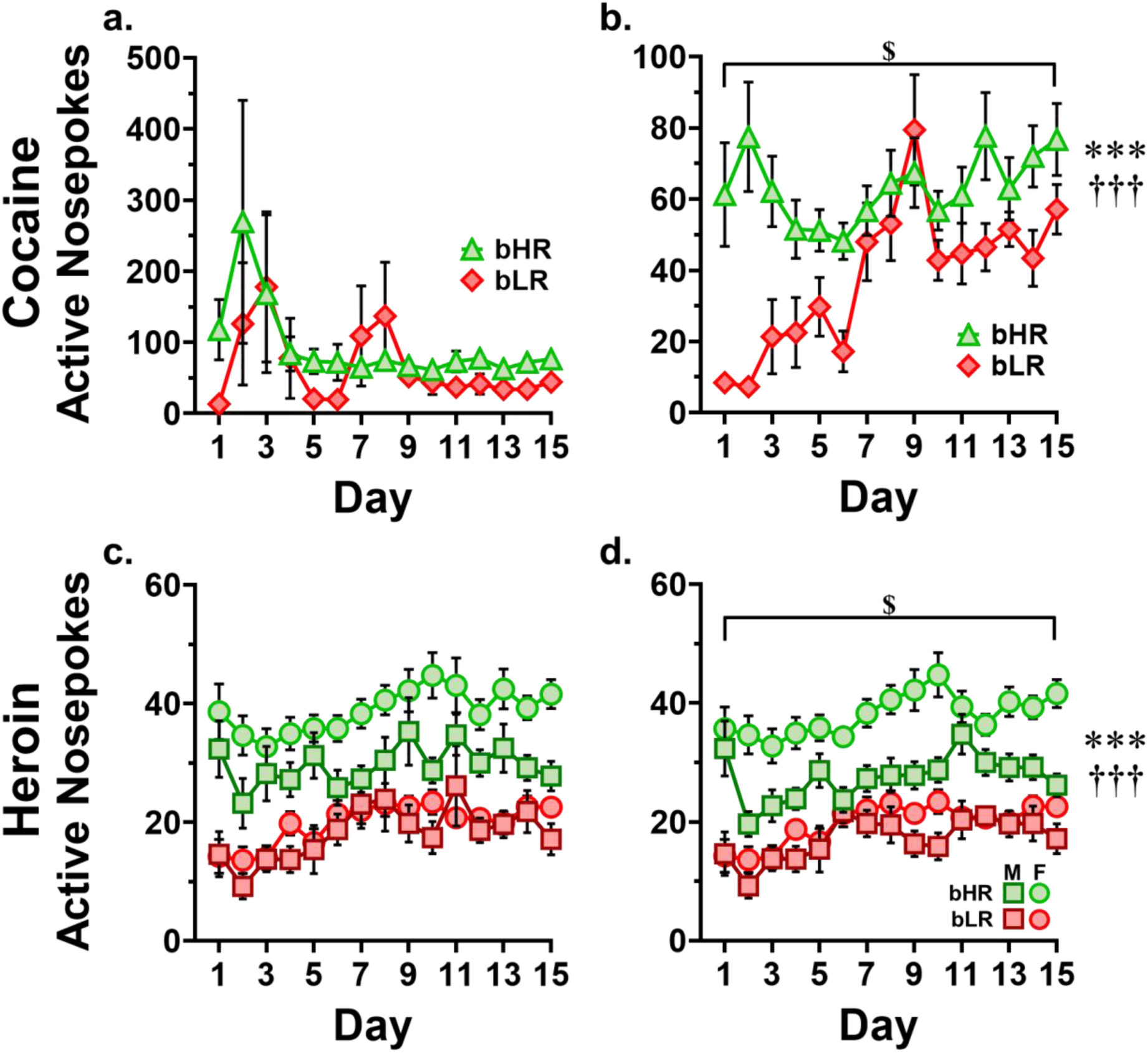
Active nosepokes (drug seeking) by day. **a.** Cocaine condition, all responders. **b.** Cocaine condition, filtered for outliers. *** line by time interaction (*p* = 0.002). ††† main effect of line (*p* < 0.001). $ main effect of time (*p* < 0.001). **c.** Heroin condition, all responders. **d.** Heroin condition, filtered for outliers. *** line by sex interaction (*p* < 0.001). ††† main effect of line (*p* < 0.001). $ main effect of time (*p* < 0.001). Box and whisker plots used to identify outliers via the Tukey method can be found in supplemental figure 1.

For heroin, the Tukey method was again used to identify and remove outliers, with the addition that outliers were identified within each line/sex subgroup, based on the finding of a sex difference in heroin infusions and therefore the likelihood of a sex difference in active nosepokes. Data presented in Figure 3c includes all collected datapoints. Data presented in Figure 3d has been processed to remove outliers. Like cocaine, Box-and-Whisker plots used to identify outliers for removal can be found in Supplemental Figure 1. There was a significant line by sex interaction (*F*_1, 388_ = 15.39, *p* < 0.001) as well as main effects of time (*F*_1, 388_ = 35.81, *p* < 0.001), line (*F*_1, 388_ = 204.39, *p* < 0.001) and sex (*F*_1, 388_ = 35.99, *p* < 0.001). The line by sex interaction was characterized by bLRs showing no sex difference in active nosepokes (*t*_193_ = 1.55, *p* = 0.123) whereas bHRs demonstrated a sex difference (*t*_199_ = 6.21, *p* < 0.001), with females showing higher active nosepoking behavior (data shown in Table 3). This correlates to what is seen in rewarded nosepokes/infusions administered.

#### 1.) Unrewarded nosepokes as a percentage of total nosepokes

Since some number of active port nosepokes will necessarily result in an infusion, a trend between active nosepokes and infusions is logical. However, this analysis fails to distinguish between a rat who has a high number of active nosepokes and a correspondingly high number of infusions, from a rat with the same high number of active nosepokes and a relatively low number of infusions, who is seeking drug at high levels when it is unavailable. A cleaner picture of the degree of drug seeking behavior is captured by asking how many active nosepokes performed in a session went *without reward*; that is, how often did the rat seek drug when it was explicitly unavailable. This was computed as the total number of active nosepokes, minus the number of infusions, divided by the number of active nosepokes, for each rat each day, representing the percentage of all active nosepokes were without reward. This allows the decoupling of high drug taking and associated nose poking from a high level of drug seeking, relative to the amount of drug taken. When this analysis is performed, a few interesting findings emerge that were not seen previously.

For cocaine, when analyzed across time, no effect of line is seen (*F*_1, 119_ = 1.282, *p* = 0.26). However, a significant main effect of time is seen (*F*_1, 119_ = 23.42, *p* < 0.001). When examining only active nosepokes that occur without reward, both bHRs and bLRs begin with high levels of drug seeking that is moderated over time until, near the end of the experimental run, both lines are rarely seeking drug when it is unavailable (Figure 4a). When examining the average percentage of unrewarded active nosepokes across the experiment, accounting for bursts of high degrees of seeking behavior, bLRs show a significantly higher percentage of unrewarded active nosepokes than bHRs (bHRs mean ± SEM 16.33% ± 1.6%, bLRs mean ± SEM 23.08% ± 2.63%, *t*_28_ = 2.192, *p* = 0.03, Figure 4b).

**Figure 4.**
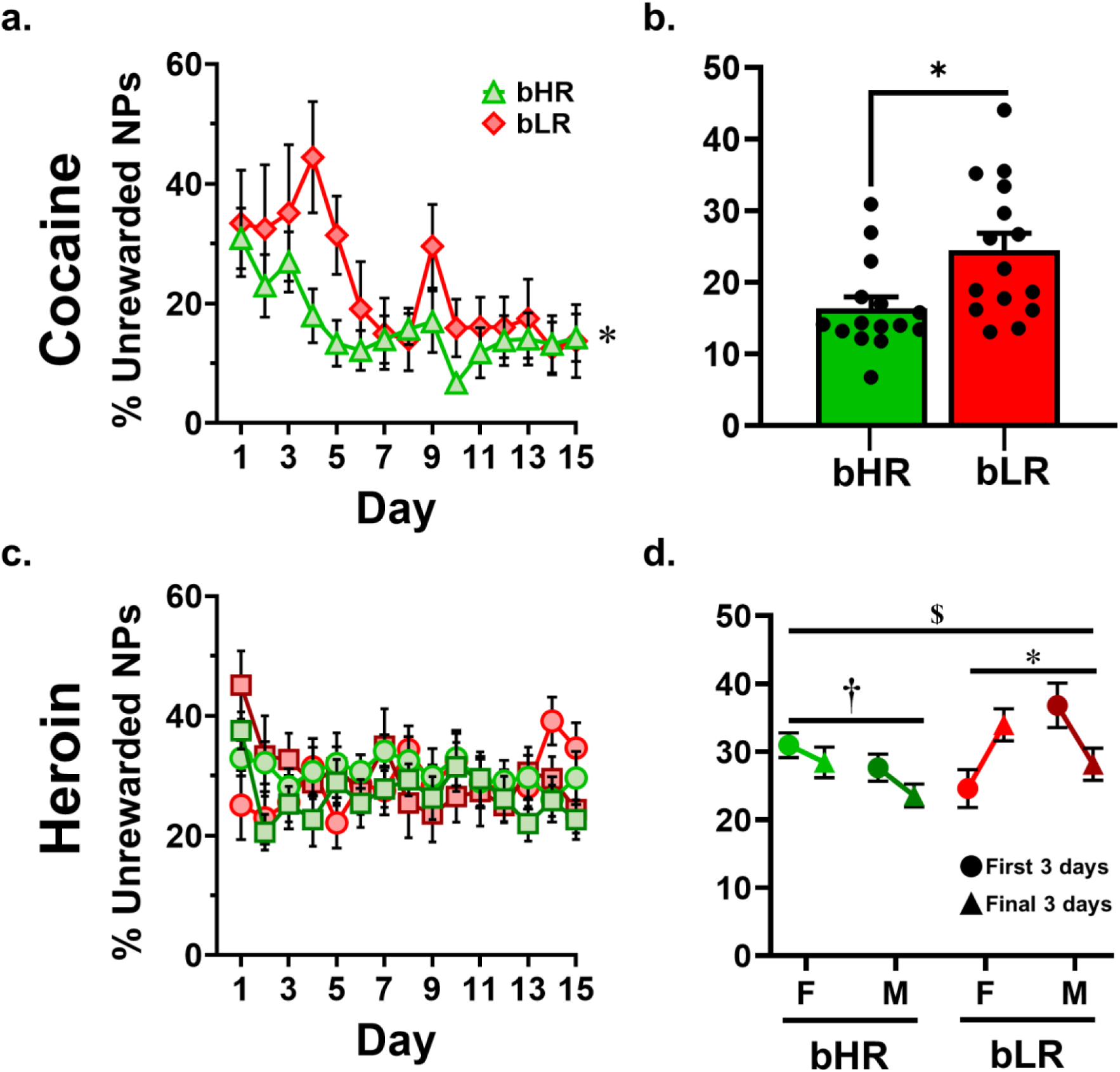
Unrewarded nosepokes as a percentage of total nosepokes. **a.** Cocaine condition, across all days. * main effect of time (*p* < 0.001). **b.** Cocaine condition, averaged across all days. * main effect of line (*p* = 0.03). Despite no line differences at the beginning or end of self-administration, bLRs demonstrate a higher overall percentage of unrewarded nosepokes, indicating a slower rate of behavioral moderation compared to bHRs. **c.** Heroin condition, across all days. Note that no group significantly moderates their percentage of unrewarded nosepokes over time when seeking heroin, compared to a significant moderation in both bHRs and bLRs when seeking cocaine. **d.** Heroin condition, examining the first 3 days vs last 3 days of behavior. While a sex difference is observed in bHRs, where males show less unrewarded nosepokes than females († *p* = 0.036), no effect of time was observed. For bLRs, a significant sex by time interaction was observed (* *p* < 0.001) where females increased and males decreased their percentage of unrewarded nosepokes over time. A line difference was also observed ($ *p* = 0.037) where bLRs had significantly higher percentages of unrewarded active nosepokes as compared to bHRs, overall.

When the heroin condition is analyzed across time (Figure 4d), a complex 3-way interaction was detected for all 3 variables (line by time by sex, *F*_1, 372_ = 7.383, *p* = 0.007) as well as an interaction of sex by time (*F*_1, 372_ = 10.22, *p* = 0.002), a line by sex interaction (*F*_1, 372_ = 5.56, *p* = 0.019), with no line by time interaction, no main effect of sex or time, but a significant main effect of line (*F*_1, 372_ = 4.38, *p* = 0.037). Follow up analysis revealed a main effect of sex in bHRs (*F*_1, 196_ = 4.46, *p* = 0.036), but no effect of time and no interaction between the two factors, indicating that in the bHR line, females had a higher overall percentage of unrewarded active nosepokes irrespective of time (females 29.7% ± 1.4%, males 25.6% ± 1.3%). Analysis of bLRs indicated a significant interaction of sex by time (*F*_1, 176_ = 12.91, *p* < 0.001) where males have more unrewarded nosepokes than females at the beginning, shifting to fewer unrewarded nosepokes and females shifting towards more unrewarded nosepokes over time. The significant main effect of line was driven by bLRs, overall, displaying greater levels of unrewarded active nosepokes (mean ± SEM = 31.1% ± 1.2%) as compared to bHRs (mean ± SEM = 27.7% ± 1.1%). Notably, there was no main effect of time for the heroin condition in any subgroup (Figure 4c).

### Inactive Nosepokes

For cocaine, inactive nosepokes did not differ based on any analyzed factor, and were expectedly low, with a median score of 1 (Figure 5c).

**Figure 5.**
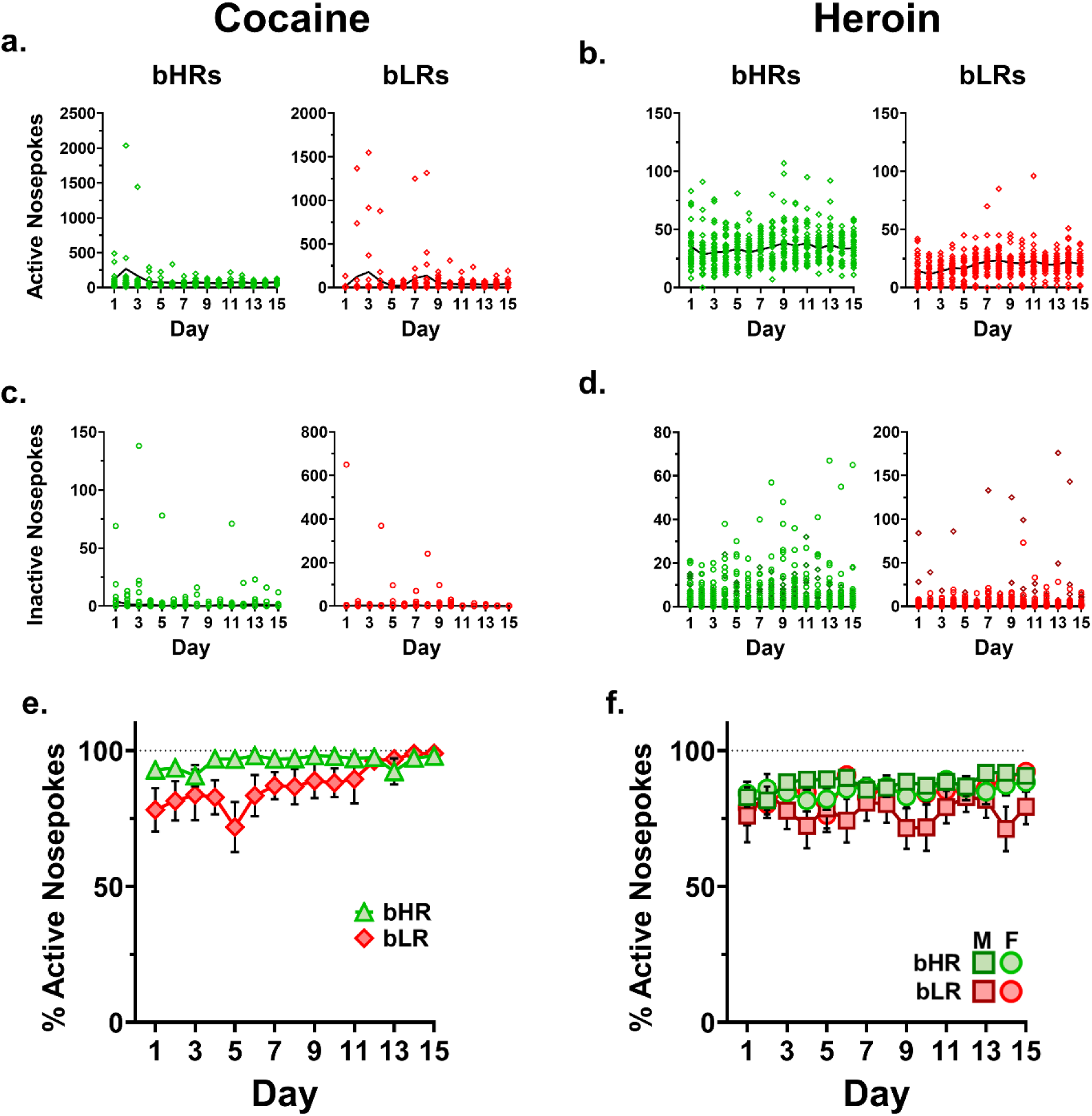
Active and inactive nosepokes by day. **a., b.** Active nosepokes, all animals, for cocaine (a.) and heroin (b.). This data is what the graphs in Figure 3, a. & c. represent, and which was analyzed via the Tukey method (Supplementary Figure 1) to identify and filter outliers. **c., d.** Inactive nosepokes, all animals, for cocaine (c.) and heroin (d.). Despite high levels of inactive nosepokes produced by some animals on some days, this does not reflect indiscriminate seeking, as the percentage of total nosepokes that are on the active port are consistently high for both cocaine (**e.**) and heroin (**f.**).

For heroin, inactive nosepokes were similarly low, with a median score of 2. However, there was a significant interaction of line & sex (*F*_1, 398_ = 12.94, *p* < 0.001), with bHR females and bLR males demonstrating higher levels of nosepoking at the inactive port than their counterparts (Figure 5d).

Despite low levels of nosepoking at the inactive port by most rats, most of the time, there were noted bursts of high levels of inactive nosepokes by individual rats. High numbers of nosepokes in the inactive port can indicate a failing catheter (and associated drug-searching by the rat), which was ruled out in this case by all rats passing methohexital catheter checks each weekend throughout the study. It can also indicate indiscriminate behavior associated with a failure to learn the drug contingency, but this was also ruled out because most rats with high inactive nosepoke counts had even higher active nosepoke counts, indicating a strong association between the active nose port and the delivery of the drug. Note that behavioral stereotypy focused on the inactive noseport cannot be ruled out. These data are displayed, for total active nosepokes, total inactive nosepokes, and the percent of all nosepokes that were at the active port (Figure 5e & 5f). There were a few animals for whom this was not the case, which will be explored at greater length in the discussion.

### Interinfusion Interval

For analysis of cocaine interinfusion interval, bLR-non-takers were excluded from analysis; as non-takers with 0 infusions taken produced an III of 120 min (the entirety of the session), significantly skewing the average III for the bLR group (Figure 6a visually displays the strength of this effect). Analysis of the average interinfusion interval, the time elapsed on average between *rewarded* active nosepokes (which due to the lockout period, can occur at a minimum of every 20 seconds) revealed a significant line by time interaction (F = 13.55, *p* < 0.001), as well as a significant main effect of both line (F = 22.40, *p* < 0.001) and time (F = 16.13, *p* < 0.001). While bHRs self-administer at a faster rate than bLRs, and both lines decrease III over time (i.e. take cocaine faster), bLRs decrease III faster over time than bHRs. These findings are summarized in Table 4 and displayed in Figure 6b.

**Figure 6.**
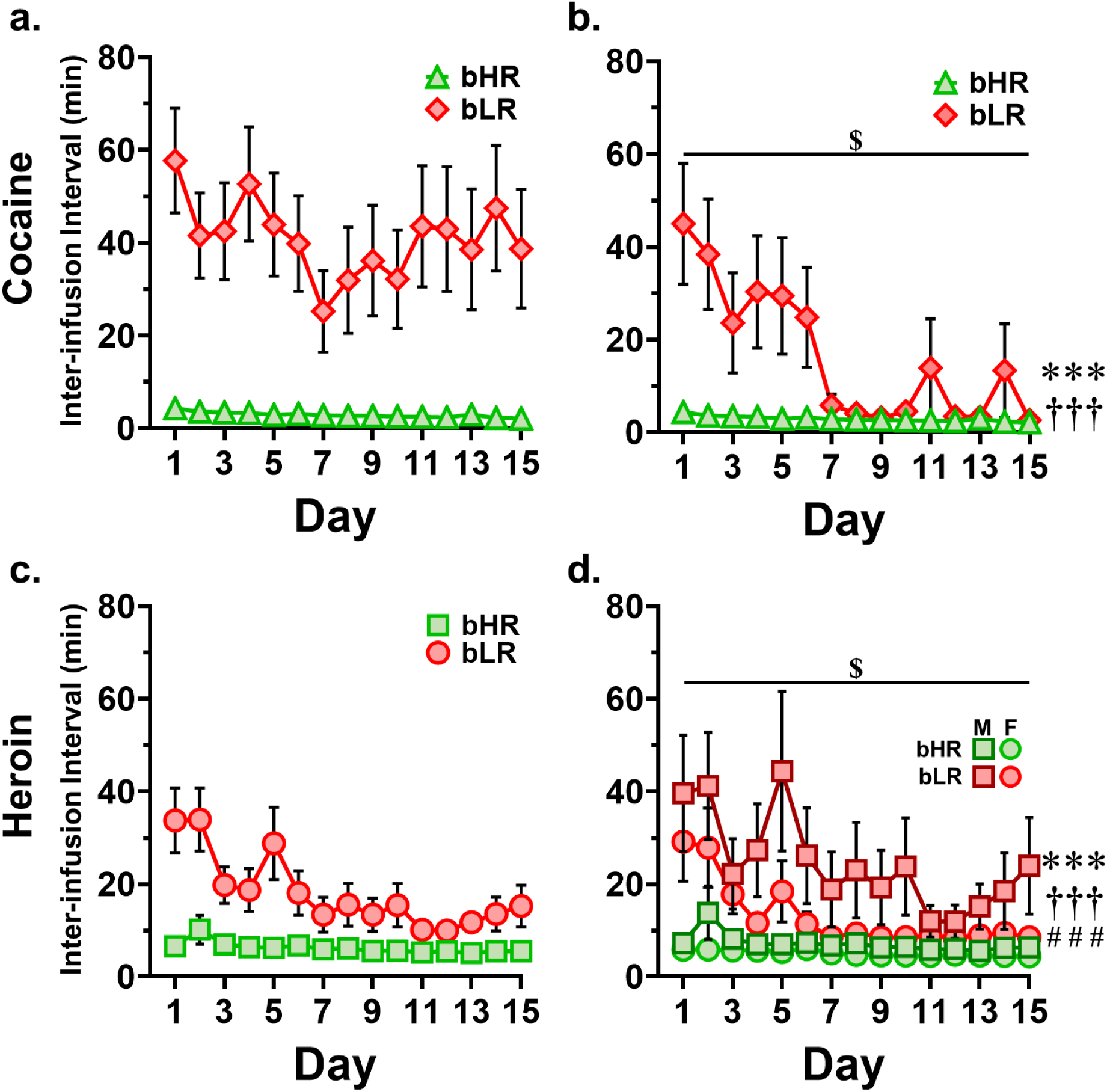
Inter-infusion interval (III) by day. **a.** Cocaine, including bLR non-responders. Due to non-responders having extremely high inter-infusion intervals, this skews the bLR group average III considerably higher as when non-responders are removed (compare to panel b). **b.** Cocaine, excluding bLR non-responders. *** line by time interaction (*p* < 0.001). ††† main effect of line (*p* < 0.001). $ main effect of time (*p* < 0.001). **c.** Heroin, with sexes combined. **d.** Heroin, with sexes separated. *** line by time interaction (*p* = 0.002). ††† main effect of line (*p* < 0.001). ### main effect of sex (< 0.001). $ main effect of time (*p* < 0.001).

**Table 4.** Summary of mean cocaine interinfusion interval, in minutes, by time and line.

| Line | Timepoint | Cocaine | Heroin |
| --- | --- | --- | --- |
| bHR | First 3 days | 3.75 (0.63) | 7.96 (1.11) |
|  | Last 3 days | 2.50 (0.26) | 5.38 (0.18) |
| bLR | First 3 days | 36.00 (6.91) | 24.00 (2.99) |
|  | Last 3 days | 6.49 (3.49) | 8.72 (0.30) |

Data are presented as mean (SEM).

For heroin, analysis of the average III revealed a significant line by time interaction (*F*_1, 398_ = 10.05, *p* = 0.002), as well as significant main effects of line (*F*_1, 372_ = 55.35, *p* < 0.001), sex (*F*_1, 372_ = 9.33, *p* = 0.002), and time (*F*_1, 372_ = 19.59, *p* < 0.001). As expected, bHRs self-administer heroin at a faster rate than bLRs, females faster than males in both lines, and self-administration rate speeds up over repeated exposures. Additionally, bLR self-administration rate speeds up relatively more over time than bHR administration rate of heroin. These findings are summarized in Table 4 and shown in Figure 6d.

## DISCUSSION

Analysis indicated that line differences existed in the drug taking behavior of both drugs, with bHRs taking significantly more drug, and at a significantly faster rate, than bLRs regardless of drug. This reflects what has consistently been seen in this model, both using outbred HR/LRs and with the selectively bred lines, for stimulants. We would like to emphasize that this is the first experiment to look at differences in opioid self-administration using a free access paradigm in either the outbred or selectively bred HR/LR model to the authors’ knowledge, and among relatively few to examine opioids in these models compared to psychostimulants. Interestingly, both lines exhibited drug-specific differences in taking behavior over the acquisition period.

Further, bLRs differ dramatically in the temporal pattern of drug taking within session as evidenced by interinfusion interval, particularly in the first week of self-administration. For instance, while the study was not designed to directly compare across the two drugs, some observations can be made regarding interinfusion interval timing in comparison to the number of infusions taken. Although the total number of infusions taken by bLRs on Day 1 of self-administration are not significantly different between drugs (heroin mean ± SEM = 8.8 ± 1.12, cocaine mean ± SEM = 10.58 ± 5.04, *t*_40_ = 0.50, *p* = 0.736), the interinfusion interval, the average time elapsing between infusions, was higher relative to bHRs for cocaine than for heroin. This indicates that bLRs take heroin significantly faster, relative to bHRs, within the first session than cocaine, despite equivalent total numbers of infusions taken.

An interesting finding to emerge from this study is the subgroup of bLRs we’ve deemed “non-takers”. While most bLR rats take cocaine at low rates early on, most will eventually accelerate cocaine taking to rates comparable to bHRs by the end of the drug acquisition phase. However, a subgroup of bLRs never accelerate cocaine taking. Regular catheter patency tests demonstrate that this is not due to a failure to adequately feel the effect of cocaine. Additionally, while some of these animals on some days take no infusions, and none ever take more than 3 infusions in a given day (see Supplemental Figure 2), all of them take some infusions most days (1.03 ± 0.91 infusions). In other words, all bLR non-takers take some cocaine, and all non-takers take cocaine more days than they do not take any cocaine. Further, these rats have an average active nosepoke percentage score of nearly 60% (57.52 ± 5.82%) across all 15 sessions. This indicates that their low level of cocaine taking is not due to failed learning of the drug delivery contingency. These rats are apparently able to feel the effects of cocaine, and seem to understand the contingency required to deliver cocaine, but never grow interested in seeking and taking it at high levels.

Multiple interesting findings emerge when the proportion of unrewarded active nosepokes is analyzed. First, strong drug-specific differences are seen, where cocaine drives a high degree of drug seeking when drug is unavailable at the beginning of the paradigm, with drug seeking becoming much less impulsive for both lines by the end of the experimental course (albeit more slowly for bLRs). Note that nosepoking when drug was unavailable did not differ by line at either the beginning or the end despite line differences in cocaine taking behavior at both timepoints, emphasizing that differences in cocaine consumption are not driven by higher impulsive or non-directed behavior in bHRs.

For heroin, there was notably no main effect of time in any group; that is to say, in stark contrast to cocaine, unrewarded active nosepoking did not significantly decrease over time for heroin in any group, despite steady acceleration of heroin consumption in all groups across time. This may indicate that heroin drives more impulsive drug seeking behavior than cocaine. Alternatively, it could indicate that heroin is a less salient incentive motivator, resulting in less potent honing of behavior over time.

Despite the lack of main effect of time in the heroin condition, there was a complex interaction of line, sex and time for heroin that was not present for cocaine. Specifically, bHR females showed modest but significantly higher levels of unrewarded seeking behavior compared to males, who were more measured in their heroin seeking behavior. The bHRs in general, regardless of sex, showed less unrewarded active nosepoking behavior than bLRs, who displayed a complex sex interaction over time, with females growing more impulsive and males growing less impulsive over repeated exposures to heroin.

Of note is the fact that for both cocaine and heroin, bLRs showed higher levels of unrewarded drug seeking when examined over the entire course of the experiment. Insofar as the percentage of active nosepokes in the absence of signaled drug availability can be interpreted as impulsive behavior, this indicates higher impulsivity for both drugs in the bLRs, despite consistently lower levels of drug consumption for both drugs. This is particularly surprising given that the LR phenotype has been consistently demonstrated to be less impulsive than the HR phenotype using more conventional behavioral tests of impulsivity, in both the outbred and selectively bred models (Stoffel and Cunningham 2008, Flagel et al. 2010b).

As mentioned in the Results, high inactive nosepoke counts were not generally regarded as indicative of a failure by the rat to learn the drug contingency, as they often had even higher active nosepoke counts, indicating a strong associative learning relationship between the active nose port and the delivery of the drug. However, there were a small number of rats which had repeatedly high inactive nosepoke counts, with low active nosepoke counts in comparison to their inactive counts. Such behavior typically implies a loss of catheter patency and drug seeking by the rat, which was ruled out by successful catheter patency testing. This led us to observe the rats behaving in real time to determine the cause. Indeed, we initially wondered if the boxes had been set up incorrectly and the rats were receiving incorrect/conflicting cues regarding active port and drug availability. Upon observation, we discovered that the rats displaying this pattern had in fact learned the drug reward contingency perfectly well; at the end of the lockout period, with the onset of drug availability cues, the rats would move to the active port and nosepoke to receive their drug infusion. Following this, they would move back to the inactive port and compulsively rub their snout, scratching themselves, against the edge of the nose port, incidentally racking up huge numbers of beam breaks at the inactive port. The reason why the inactive port, as opposed to the active port, became the target of this stereotyped behavior for multiple rats remains a mystery. However, behavioral stereotypies in rodents caused by drugs of abuse are common, particularly for psychostimulants. These incidents lead us to advise caution in using data such as active vs inactive port counts as an indicator of strength of the association between drug associated cues and drug reward.

One of the most consistent findings in our self-administration behaviors is a sex difference, often specific to the bHR phenotype, specifically in heroin self-administration behaviors, which is not observed in cocaine self-administration. There is a sex difference specific to bHRs in total infusions administered, active nosepokes, and inactive nosepokes, driven by females demonstrating higher behavior in every case; and a sex difference across both lines in interinfusion interval, again driven by the females. Sex differences were much less robust in cocaine self-administration behavior, which showed only a sex by line interaction in total active nosepokes where the line difference was magnified in females relative to males (but with no sex difference present within either line), which disappeared when examining only unrewarded active nosepokes.

These findings reflect an effect that is commonly seen in drug addiction studies, both in humans and rodents. Sex differences are not always reliably seen, but when they are present, they are nearly always in the direction of worse outcomes for females (for thorough review of this literature, please see Becker and Hu (2008), Becker (2016), Becker et al. (2017), and of suspected biological mechanisms underlying this effect (Hu et al. 2004)). However, it is interesting that in this study, it appears most often in a line-specific manner, specific to the bHR females, but only rarely seen in bLR females. Interestingly, a previously observed finding in much earlier generations of this bred rat model which showed significantly increased cocaine self-administration specific to bHR females and absent in bLRs (Davis et al. 2008). Note that in the current study, no sex difference was seen in cocaine infusions as was seen by Davis et al., but the sex difference in heroin self-administration was also specific to bHR females. This also reflects prior observations of other behaviors within this bred model that have been established to partially predict susceptibility to drug addiction, such as total time spent in the open arms of elevated plus maze, as well as locomotor response to a novel environment, the behavioral output used to characterize the model (Hebda-Bauer 2017). In both cases, bHR females show a heightened level of these addiction susceptibility-related behaviors (Hebda-Bauer 2017). These findings could indicate that the genetic susceptibility to addiction conferred by female sex and that conferred by bHR phenotype combine, resulting in a greater vulnerability to opioid addiction than psychostimulant addiction. It has been demonstrated that female rats accelerate opioid self-administration faster than males (Towers et al. 2024) as well as cocaine (Lynch and Carroll 1999, Lynch and Carroll 2000); thus, the sex difference found in the current study is not as remarkable in itself, as much as its strong relation with temperamental phenotype.

It should be noted that these are not only different drugs but entirely different classes of drug (psychostimulant vs opioid) and are believed to differentially drive the reward circuitry that underlies self-administration behavior, with cocaine directly increasing dopaminergic tone in the reward circuitry, particularly in striato-accumbens, and heroin indirectly increasing it via ascending disinhibition of dopaminergic circuits, as well as suspected direct opioidergic effects operating independently of dopamine (discussed further below). Because of this, comparisons between the doses of drug administered should be done with caution, despite driving similar initial levels of consumption in bLRs.

However, it has been demonstrated previously that line-specific differences in self-administration of cocaine are not due to differential potency of the drug in each line but to differential maximal efficacy; that is to say, HR and LR rats demonstrate peak locomotor response to cocaine at the same dose despite large differences in the magnitude of that peak response (Piazza et al. 2000). It is not unreasonable to assume the same is true for heroin, and locomotor responses to morphine have been shown to differ between bHR and bLR for at least one dose (Deroche et al. 1993). Thus, line-specific differences in cocaine and heroin taking are likely to be fundamentally rooted in biological differences between bHRs and bLRs as opposed to different pharmacokinetics. For example, bHRs and bLRs have different basal dopamine levels in the NAcc (higher in bHRs) and different levels of cocaine-induced dopamine release (higher in bHRs) which are likely to at least partially account for differences in cocaine self-administration behavior (Mabrouk et al. 2018). Conversely, differential preference for heroin could be at least partially accounted for by differential expression levels of both mu and kappa opioid receptors in key reward regions including the NAcc (bHRs have higher mu receptor and lower kappa receptor gene expression in the NAcc than bLRs) (Turner et al. 2019).

It remains possible that drug-specific differences in administration within each line may be due to differential relative position of the administered dose on the dose-response curve (i.e. heroin dose being closer to the peak of the dose-response curve than cocaine dose). This is impossible to determine without a full dose-response study of both drugs in both lines, which is beyond the scope of the current work. However, prior work by Piazza et al. (2000) demonstrated that the peak of the dose-response curve for cocaine was at equivalent doses for outbred HRs and LRs and that levels of cocaine in the brain were equal, and cleared along an equal timeframe, in HRs and LRs. This does not rule out the possibility that this does not hold true for opioids, but we feel this to be unlikely, particularly given prior findings regarding HR/LR locomotor response to other opioids, such as morphine (Deroche et al. 1993). Nevertheless, future work should examine the pharmacokinetics of opioids in the bHR/bLR lines for horizontal shifts of the dose-response curve between lines.

The idea that genetically mediated vulnerability to addiction could be labile and affected by the specific drug is not a new one. Research in the personality traits of substance users has revealed that personality traits related to both internalizing and externalizing temperament are somewhat predictive of drug choice/preference (Hopwood et al. 2008). It has been observed in adolescents that internalizing behaviors in the absence of externalizing problems appear to exert a protective effect against substance use problems, while internalizing problems comorbid with externalizing problems may *increase* risk of substance use compared to presence of externalizing traits alone (Scalco et al. 2014). Early research into the role of personality in drug choice found that psychostimulant users were high on Experience Seeking, Disinhibition, and Boredom Susceptibility, and tended to be highly energized individuals at baseline (Carrol and Zuckerman 1977). This parallels what is seen in bHR rats which display high baseline levels of energy and behavioral activity, high impulsivity (indicative of disinhibition) and high sensation seeking (reviewed in Flagel et al. (2014)). In contrast, opiate (and sedative) users showed low Experience Seeking, an externalizing trait, concluding that these users typically do not use drugs to seek novel sensations but rather to dull unwanted, excessive stimulation (Carrol and Zuckerman 1977). Individuals with SUD who have comorbid internalizing disorders also appear to commonly use drugs to self-medicate symptoms of internalizing disorders (Brady et al. 2004, Crum et al. 2013).

Thus, differences between bHRs and bLRs of drug seeking and taking for opiates relative to stimulants, observed previously in outbred rats (Chang et al. 2022) and in the current work in the selectively bred model, likely represents a fundamental, genetically mediated trait intrinsic of the model. The previously observed differences in both basal tone and cocaine-evoked release of monoamines, particularly dopamine, in the striatum (Mabrouk et al. 2018) may account for the differences between bHRs and bLRs in their response to, and seeking of, cocaine. However, in various human studies, opioids routinely fail to evoke a robust dopamine response (Nutt et al. 2015) and when a dopaminergic response to opioids is observed, it appears to be either uncorrelated or *inversely* correlated to hedonic experience (Spagnolo et al. 2019). Further, oxycodone and morphine have been demonstrated to drive very different dopamine release kinetics in the nucleus accumbens (Vander Weele et al. 2014) despite both drugs being readily self-administered (Ree et al. 1978, Beardsley et al. 2004, Zhang et al. 2014) and driving comparable conditioned place preference (Narita et al. 2008, Niikura et al. 2013), commonly considered a means of measuring drug reward (as distinct from reinforcement) (Bardo and Bevins 2000) Therefore, the strong differences in striatal dopamine tone between bHR and bLR rats is perhaps unconnected to seeking and taking of opioids, which may be driven by separate neurobiological differences altogether.

Recent work by Badiani and colleagues has demonstrated that in both humans and rodents, volitional drug choice between heroin and cocaine is largely dependent on environmental context (i.e. ‘home’ vs ‘outside the home’), with humans and rodents alike demonstrating stronger ‘liking’ and ‘wanting’ of heroin at ‘home’ and of cocaine ‘away’ (Badiani et al. 2019), and with rats self-administering the two compounds according to similar contextual patterns when provided the option to do so, regardless of drug history (De Luca et al. 2019). These findings further reinforce the idea that psychostimulants and opioids cannot simply be thought of as two variants or subclasses of drugs of abuse acting in broadly similar ways, but rather display a complex interaction with the environment and the individual to scaffold volitional drug taking and the transition to addiction more strongly in some individuals and circumstances and not in others. A thorough understanding of the impact of such individual differences and environmental factors may provide useful tools for the development of treatment and prevention strategies for drug abuse and addiction. Further, while we did not familiarize the animals with the self-administration chambers to differing degrees as was done by Badiani (De Luca et al. 2019), we may speculate that differential approach to cocaine and heroin by bHR and bLR rats, at least initially, may partially reflect differing perceptions about the nature of the testing environment.

Key to interpreting these results is the concept that the HR phenotype represents the highest tail of the normal rodent population from which it is drawn in its predilection to seek and take addictive substances. Indeed, it was this high motivation to seek and take drugs, relative to the population average, that the model was originally designed to predict (Piazza et al. 1989, Piazza et al. 1990). The selectively bred bHR line is likely to represent a subset of rats even more highly enriched for drug-taking, as the founding generations of the line used a particularly stringent definition of ‘high responder’ to include only the top 20% of the population (Stead et al. 2006), while many outbred screens use top/bottom quartile, top/bottom third, or even the very liberal median-split procedure to characterize HRs vs LRs. Thus, addiction-related behaviors of bLRs should not be interpreted independently, but in comparison to what the highly addiction-susceptible bHRs are doing in the same circumstances. While bLRs typically remain a statistically distinct population relative to bHRs, their behaviors shift far more bHR-like following repeated exposures to cocaine, reflecting prior findings that bLRs will acquire addiction-like behavior given adequate levels of drug exposure (Flagel et al. 2016). In comparison, bLR behavior remains distinct from bHR behavior throughout varying measures of addiction-like behavior, implying a qualitatively different trajectory of experience, and likely neurobiological impact, when exposed to opioids as compared to psychostimulants. Future work should seek to confirm that this generalizes to other opioids and is not specific to heroin, as well as investigating the differential neurobiological impacts of repeated opioid exposure relative to psychostimulant exposure.

Although the current finding may not be directly reflective of findings in humans, where internalizing traits in the absence of comorbid externalizing traits may be protective (irrespective of drug choice; but note that this body of literature is relatively sparse), it represents important evidence in support of genetically determined influences of personality/temperament on drug choice/preference. Notably, the present finding, coupled with the recent report of a complete lack of remifentanil self-administration differences in outbred HR/LR rats under a controlled dosage paradigm (Chang et al. 2022) and prior evidence of stress-induced shifts in LR stimulant taking (Kabbaj et al. 2001, Kabbaj et al. 2002), challenges the long-held interpretation that ‘only HRs are predisposed to drug-taking’ (Dellu et al. 1996) and the idea that HRs represent a ‘vulnerable’ and LRs a ‘resistant’ phenotype with regard to drug intake (Piazza et al. 2000). Instead, it is becoming increasingly clear that the LR phenotype represents a more labile form of susceptibility, dormant at baseline and especially regarding psychostimulants but able to be activated by the experience of stress, and perhaps more susceptible even at baseline when offered opioids rather than stimulants.

## ACKNOWLEDGEMENTS

This work was supported by NIDA U01DA043098; NIDA T32DA07268; Office of Naval Research (ONR) 00014-19-1-2149; The Hope for Depression Research Foundation (HDRF); and The Pritzker Neuropsychiatric Research Consortium. The authors would like to thank the University of Michigan Consulting for Statistics, Computing and Analytics Research (CSCAR) office for technical advice regarding statistical modeling, Pam Maras, Megan Hagenauer, and Huzefa Khalil for valuable advice and conversation, and Sara Westbrook for technical advice.

## AUTHORS’ CONTRIBUTIONS

Conceptualization: HA

Data curation: MAE

Formal analysis: MAE

Funding acquisition: HA

Investigation: MAE, AP, SK, EKH, KL, BDL, SC

Methodology: HA, JB, SBF, SJW

Project administration: HA

Resources: HA, SBF, JB, SJW

Supervision: HA

Visualization: MAE

Writing – original draft: MAE (lead), HA (supporting)

Writing – review & editing: MAE, HA, SBF, JB, SJW

