## Supplemental Information for "Differential acquisition of cocaine and heroin self-administration in a rat model of internalizing versus externalizing temperament"

**Supplemental Data**

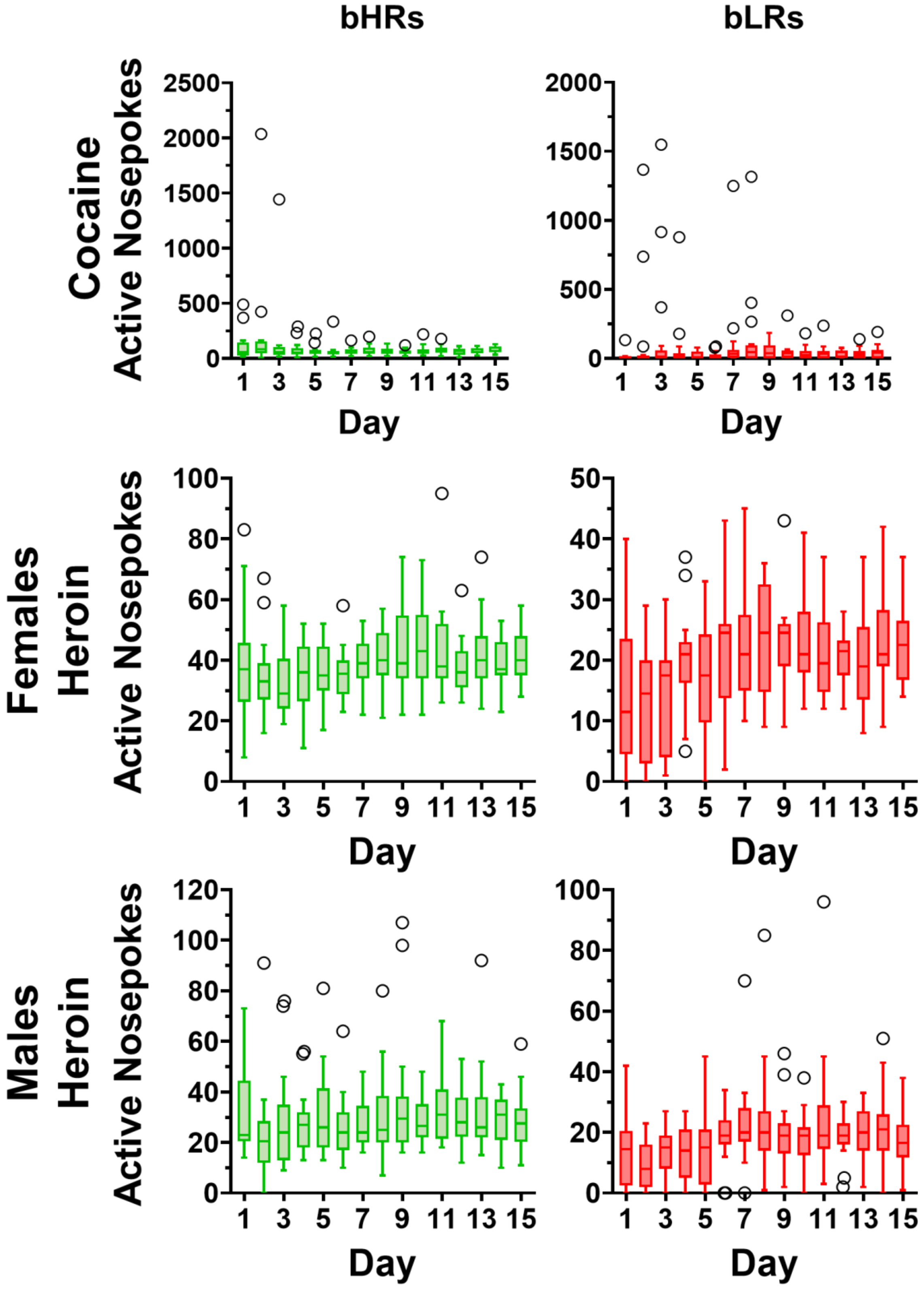

**Supplemental Figure 1. Box-and-Whisker plots used to identify outliers in active nosepoke data.** Box-and-whisker plots were created using the Tukey method, with outlier data points in black. Figure 3a and 3c include all datapoints. Figure 3b and 3d exclude the outlier data points identified here.

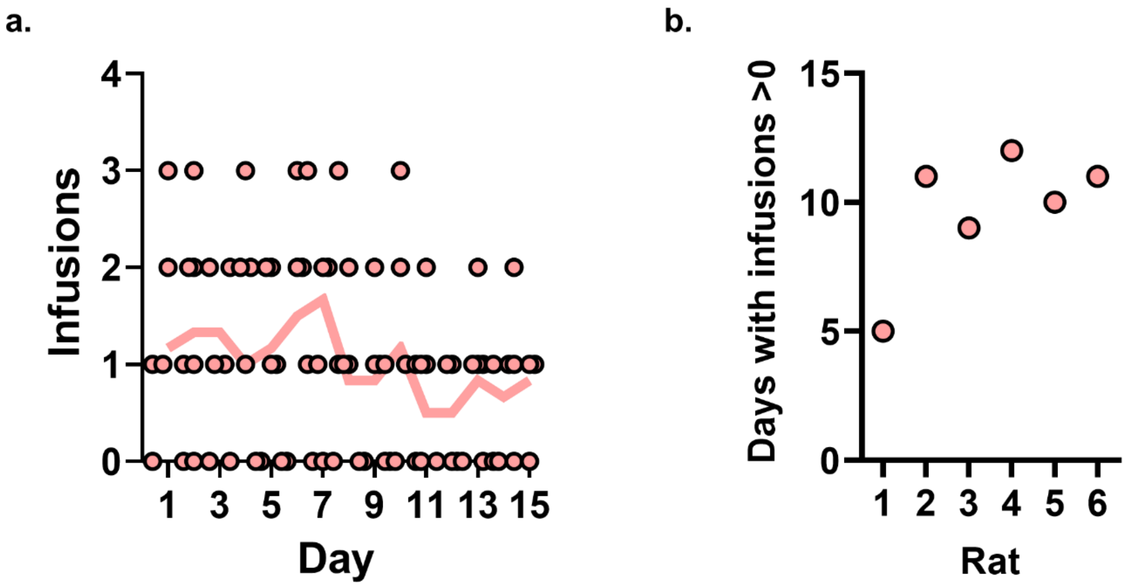


**Supplemental Figure 2. Cocaine infusions taken by bLR non-takers.** While no rat identified as a non-taker took more than 3 infusions on any given day, all rats took some infusions, and the majority took cocaine on a majority of days (b), with the daily mean number of infusions hovering near 1 (represented by the line in panel a).
